# Topological Scoring of Protein Interaction Networks

**DOI:** 10.1101/438408

**Authors:** Mihaela E. Sardiu, Joshua M. Gilmore, Brad D. Groppe, Arnob Dutta, Laurence Florens, Michael P. Washburn

## Abstract

It remains a significant challenge to define individual protein associations within networks where an individual protein can directly interact with other proteins and/or be part of large complexes, which contain functional modules. Here we demonstrate the topological scoring (TopS) algorithm for the analysis of quantitative proteomic analyses of affinity purifications. Data is analyzed in a parallel fashion where a bait protein is scored in an individual affinity purification by aggregating information from the entire dataset. A broad range of scores is obtained which indicate the enrichment of an individual protein in every bait protein analyzed. TopS was applied to interaction networks derived from human DNA repair proteins and yeast chromatin remodeling complexes. TopS captured direct protein interactions and modules within complexes. TopS is a rapid method for the efficient and informative computational analysis of datasets, is complementary to existing analysis pipelines, and provides new insights into protein interaction networks.

## Introduction

Many large and medium scale analyses of protein interaction networks exist for the study of protein complexes ^1–4^. These studies typically employ a quantitative proteomic analysis of affinity purifications analyzed using mass spectrometry (APMS) of bait proteins and utilize a statistical tool to provide a confidence that a prey protein is associated with a bait protein. Approaches like COMPASS ^5^, QSPEC ^6^, SAINT ^7^, and SFINX ^8^ largely yield statistical values, like a *p* value, to provide a confidence that two proteins are associating or are part of a protein complex.

Within a protein interaction network, an individual protein may have multiple interactions, may be part of a large protein complex or complexes, and large protein complexes can be composed of important functional modules. For example, modularity is a hallmark of protein complexes involved in transcription and chromatin remodeling. Within this area of a protein interaction network Mediator ^9^, SAGA ^10, 11^, SWI/SNF ^12^ are but a few of the many complexes well known to have modules that carry out distinct functions. Determining these modules in these complexes has required years of study using biochemical, genetic, and proteomic methods ^9–12^. In addition, within protein interaction networks and protein complexes there are also direct protein interactions that are critical for biological functions. For example, in DNA repair, the Ku70-Ku80 (XRCC5-XRCC6) heterodimer is critical for recognition of DNA double strand breaks that occur during non-homologous end joining ^13^. Existing statistical tools struggle to gain insight into the define the behavior of an individual protein in a protein interaction network remains a major challenge. New methods are needed to gain deeper insight into protein interaction networks and determine direct protein interactions and capturing modularity in large network dataset.

We have used approaches like deletion network analyses, network perturbation, and topological data analysis to determine the modularity in protein interaction networks and protein complexes themselves ^14^. Here we describe a novel topological scoring (TopS) algorithm for the analysis of quantitative proteomic APMS datasets and protein interaction networks. TopS can be used by itself or in addition to existing tools ^5–8^ to analyze and interpret datasets. Here we applied TopS to a human DNA repair based protein interaction network dataset and previously published yeast chromatin remodeling datasets from the INO80 and SWI/SNF complexes ^12, 14^. TopS yielded insights into the direction protein interactions and modularity within these networks. TopS is a simple and powerful method of inferring the interaction preferences of proteins within a network consisting of reciprocal and non-reciprocal purifications based on the likelihood ratio method. The TopS algorithm generates positive or negative values across a broad range for each prey and bait relative to the other baits APMS analyses in a dataset. TopS can differentiate between high-confidence interactions found with large positive values and lower confidence interactions found with negative values. TopS has the advantage that the values themselves can be easily integrated into additional computational workflows and clustering approaches.

## Results

### Definition of Topological Score Based on Likelihood Ratio across Aggregated APMS Datasets

Determining whether a protein in a single affinity purification is significantly enriched without additional information remains a challenge. Therefore, by adding other biologically-related baits to the dataset, one can determine whether a protein is truly enriched in a sample. The concept behind computing TopS is to be able to collectively analyze parallel proteomics datasets and highlight enriched interactions in each bait relative to the other baits in a larger biological context. For each individual bait, instead of calculating a score by concentrating only on a single bait column via normalization or modelling, we now aggregate information from the whole dataset where all data from all rows and all columns are used. Our topological score is based on the likelihood ratio and reflects the interaction preference of a prey protein in a bait. For each protein detected in a bait APMS, TopS calculates the likelihood ratio between the observed spectral count *Qij* of a protein *i* in a bait *j*, and the expected spectral count *Eij* in row *i* and column *j* (Figure 1). Each bait protein from each APMS experiment has a unique TopS value. TopS assigns positive or negative scores to proteins identified in each bait using spectral count information. If the actual number of spectra of a prey protein in a specific bait APMS exceeds that in all baits APMS, the presumption is that we have a positive preferential interaction. Likewise, fewer spectra in the bait APMS indicate a negative interaction preference.

**Figure 1.**
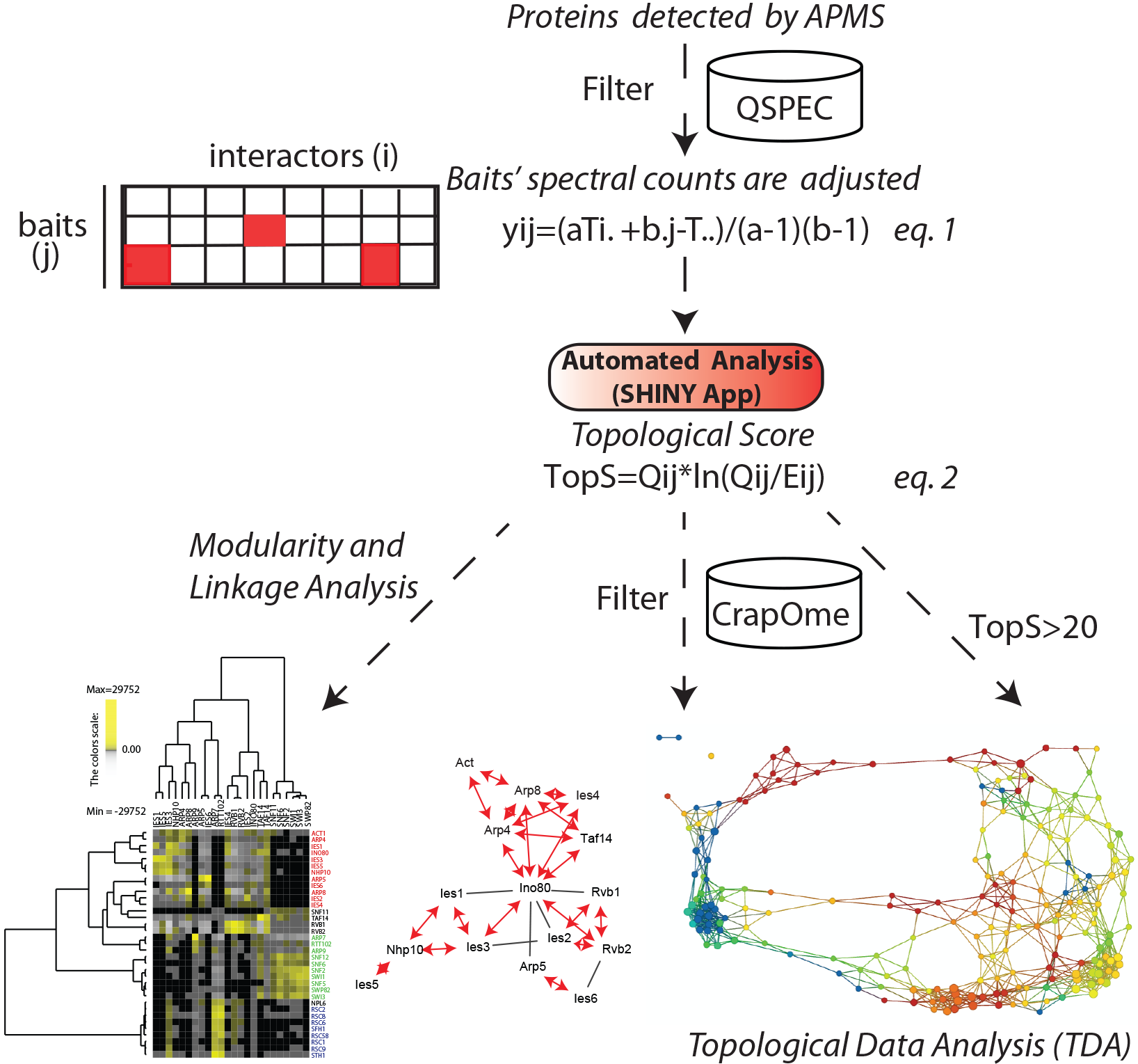
Workflow for Topological Scoring of Protein Interaction Datasets. In the first step, contaminant proteins are filtered using Z-scores and FDR generated from QSPEC, however other approaches could be used. Next, overexpressed bait proteins that have high spectral counts in the data are adjusted using equation 1. After prey proteins as baits are adjusted, topological scores (TopS) are then calculated, using an automated SHINY application, as described in equation 2, where Qij is the observed count in row i and column j and Eij is the expected count in row i and column j. Direct input of TopS values can be used in many different ways, such as data clustering to investigate the modularity and linkage in a network; additional filtering may be conducted using the CrapOme, for example; and a topological data analysis network may be generated.

Unlike *p*-values or fold changes where the difference between the largest and the smallest values is relatively small, TopS generates a wide range of positive and negative scores that can easily differentiate high, medium, or low interaction preferences within the data. This is an advantage for proteomics data analysis since these scores not only reflect the interaction preference of proteins relative to others, but can now be directly integrated to further analyses such as clustering or network analysis in order to discover network organization. TopS is written in R and the platform is built with SHINY (https://shiny.rstudio.com/). It is easily applicable and it includes correlations and clustering for bait relationships.

### Analysis of a Transiently Overexpressed Human DNA Repair Network Dataset

We first tested TopS on a human DNA repair protein interaction network dataset generated from HaloTag™ proteins transiently expressed in HEK293 cells. DNA repair mechanisms are complex, independent, yet have extensive crosstalk, and have been extensively studied ^15, 16^. To uncover the connectivity between proteins involved in these pathways, we selected 17 proteins that are part of different DNA repair mechanisms. In addition to the affinity purifications of known elements of the DNA repair pathways such as MSH2, MSH3, and MSH6 (involved in mismatch repair),RPA1, RPA2, and RPA3 (involved in nucleotide excision repair), XRCC5 and XRCC6 (involved in double strand break repair), SSBP1 and PARP1 ^17^, we analyzed an additional 7 proteins (WDR76, SPIN1, CBX1, CBX3, CBX5, CBX7, and CBX8) with chromatin associated functions and some of which are associated with DNA repair ^18^. For example, we have previously demonstrated that WDR76 is a novel DNA damage response protein with strong associations with members of DNA repair pathways and the CBX proteins ^19^. It is important to note that our objective here was not to describe a human DNA repair protein interaction network. The proteins chosen for this small scale study were of interest because of their potential relationship to the poorly characterized WDR76 protein ^19^. Also, it is important to note that these were transiently overexpressed proteins in HEK293 cells, and given the potential issues with transient overexpression ^20^, the dataset would be expected to be noisy. We reasoned that using a noisy dataset would be an excellent test of the TopS method and its ability to extract meaningful biological information.

We used Halo affinity purification followed by quantitative proteomics analysis to identify proteins associated with either one of the 17 baits. Three biological replicates were performed for each of the bait proteins and the distributed normalized spectral abundance factor (dNSAF) ^21^ were used to quantify the prey proteins in each bait. To eliminate the potential non-specific proteins, three negative controls were performed from cells expressing the Halo tag alone. A total of 54 purifications were completed and 4509 prey proteins identified (Supplemental Table 1). To determine the proteins that were enriched in the samples versus negative controls, the QSPEC ^6^ application was used (Supplementary Table 1). A protein was considered specific in a sample if the Z-score was greater than or equal to 2 and the FDR was less than 0.01 in the bait APMS versus the control APMS. A total of 801 prey proteins passed these strict statistical criteria and they were further used in the analysis. Because of the higher number of spectra identified for overexpressed bait proteins, we adjusted the spectral counts for each bait protein according to *equation 1* (Figure 1 and Methods). To depict the interactions in this DNA repair dataset that consisted of a matrix of 801 proteins in 17 baits, we calculated topological scores based on equation 2 and assigned positive or negative TopS to proteins identified in each bait APMS. To focus on proteins with positive interaction preferences, we used a TopS cutoff of 20 (Supplementary Fig. 1). A total number of 617 proteins passed this filtering criteria (Supplementary Table 2).

Next, we investigated whether TopS values could be used directly for clustering and to determine if baits were properly separated. In addition, we hypothesized that proteins within the same complex should have high preferences to same bait purification. Therefore, we separated proteins in complexes using ConsensusPathDB ^22^ and systematically examined their TopS values. We detected 118 protein complexes consisting of 230 proteins. It is important to note that some proteins were shared by multiple protein complexes. These results demonstrated that proteins within complexes tend to associate with the same baits with high topological scores (Supplementary Table 3 and Fig. 2) even though some of the proteins were detected in most of the baits. Thus, by using TopS values, we could illustrate the preferential interactions between protein complexes and baits in a large dataset. To further investigate this observation, we clustered all of the proteins in the detected complexes and we obtained a strong separation of the baits demonstrating that the topological scores are indeed significant (Fig. 2A). In addition, bait proteins that belong to the same complexes clustered together (Fig. 2A).

**Figure 2.**
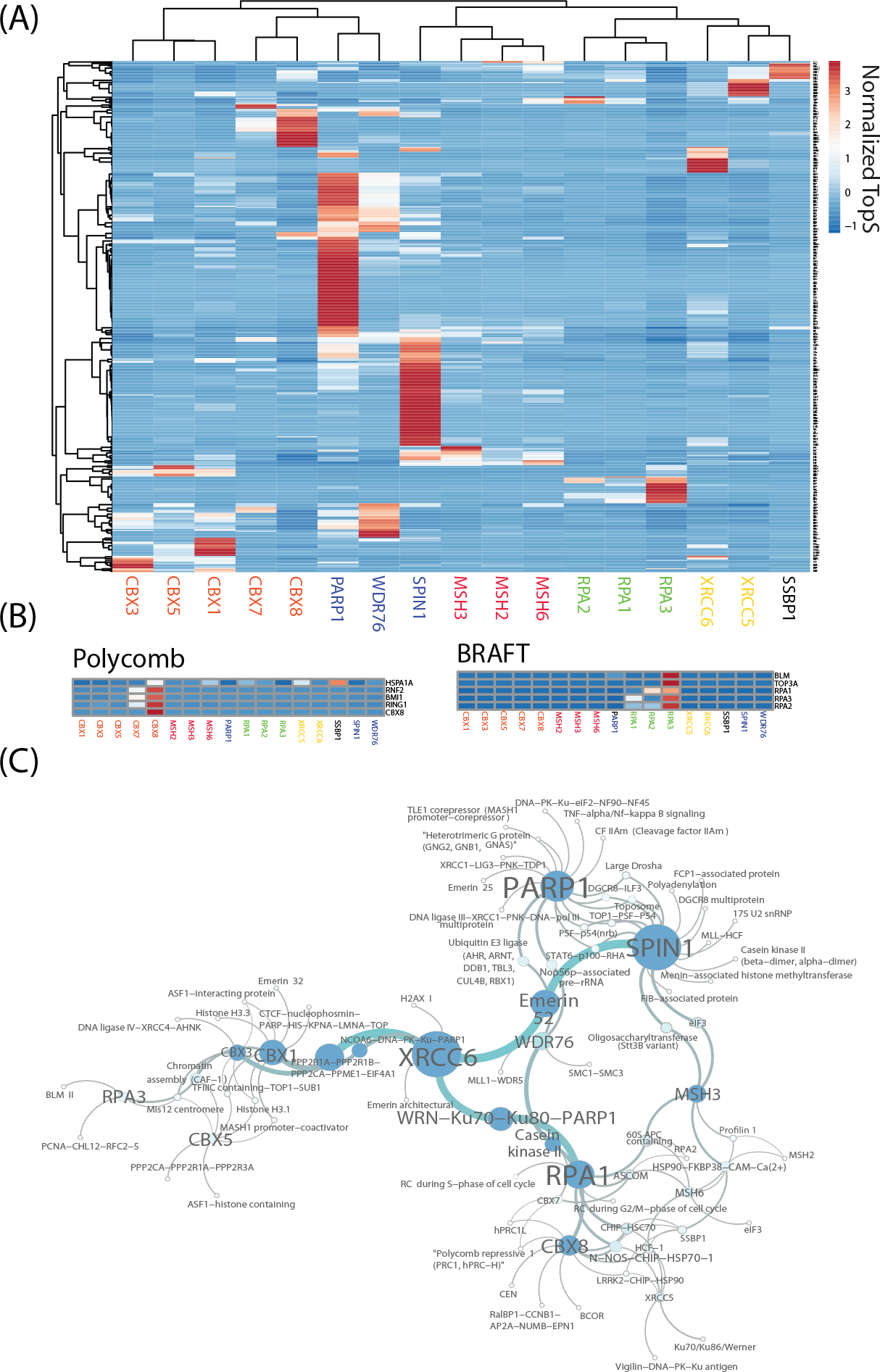
Capture of Protein Complexes in a Human DNA Repair Network. (A) Hierarchical biclustering on TopS values of the 17 baits and 535 preys in a human DNA repair protein interaction network. Proteins identified in complexes in the DNA repair dataset were hierarchically clustered using normalized TopS values as input using ClustVis ^39^. A scaling method was used to divide the values by standard deviation so that each row had a variance equal to one. Rows were centered and unit variance scaling was applied to rows. Rows were clustered using correlation distance and average linkage. Columns were clustered using correlation distance and Ward linkage. (B) Bait-specific protein complex enrichment. Two complexes are shown that showed high association with specific baits. Red color corresponds to high TopS scores. Subunits of the polycomb complex showed high scores with the CBX8 bait, and components of the BRAFT complex showed high scores with the RPA3 bait. (C) Interaction network between baits and known protein complexes identified in the dataset. The network was constructed using the Cytoscape platform ^23^. Large nodes correspond to a larger number of links.

Certain complexes showed significant enrichment to some of the baits. For example, in the case of the polycomb complex, we detected 5 members of the complex including CBX8. These five proteins had high preferential interactions to the CBX8 bait (Fig 2B). Similar results were observed in the case of the BRAFT complex where we observed that five components of the complex, including RPA proteins, show high scores in the RPA3 bait APMS dataset confirming the connectivity between these proteins (Fig 2B). Next, to further illustrate all of the connections between complexes and baits we next constructed a network using the Cytoscape platform ^23^ (Fig 2C). This network illustrates that PARP1, SPIN1, and XRCC6 baits have the most connections, and PARP1 and SPIN1 share a significant connected subnetwork (Fig 2C).

### Topological Network Assembly and Extreme Value Analysis in the Human DNA Repair Network

Next, we assembled a protein interaction network based on Topological Data Analysis (TDA) ^24^ using TopS values (Fig. 3A). We previously demonstrated that TDA ^24^ can be used to identify clusters of proteins in protein interaction networks using normalized correlations ^25^ and to find topological network modules in perturbed protein interaction networks using fold change ratios ^14^. TDA in conjunction with TopS values resulted in proteins with high topological scores to cluster to the same region of the network. For example, proteins that are likely to interact with PARP1 are located on the right side of Figure 3 in the same topological area, whereas proteins that interact with WDR76 are in the upper left side of Figure 3 (Supplementary Table 4). WDR76 is in the same node with another 18 proteins including HELLS, GAN, and SIRT1, which were also observed by independent studies to associate with WDR76 ^2, 26^. Similarly, SPIN1 is in a node with another 18 proteins, four of which (SPIN1, SPIN4, THRAP3 and BCLAF1) have been reported by others in SPIN1 purifications ^2^. Additional areas are highlighted to show associations with SPIN1, CBX proteins, and XRCC5, for example (Fig. 3A).

**Figure 3.**
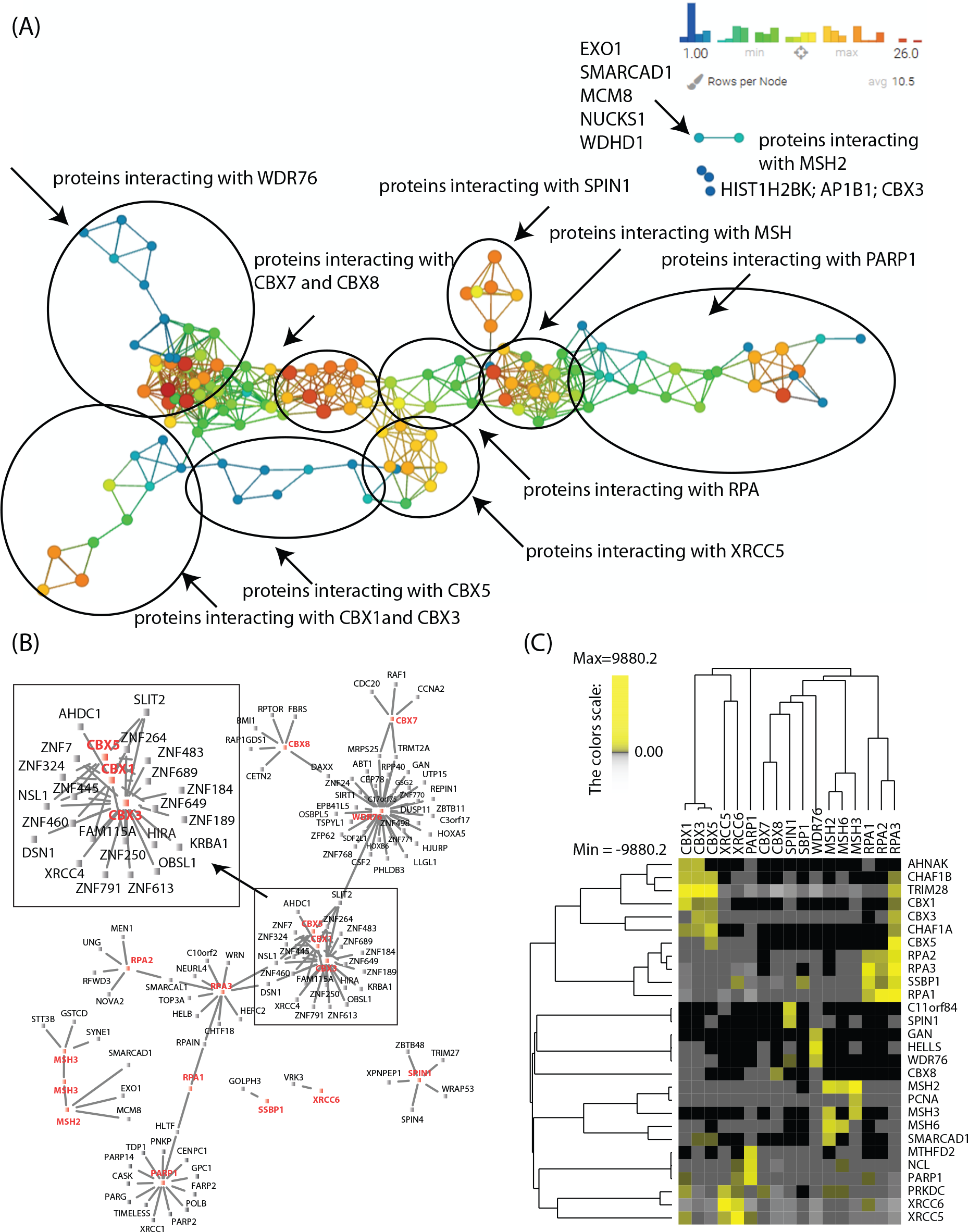
Topological Network and Linkage Analysis of Human DNA Repair Network. (A) TDA in combination with the TopS scores was applied to the proteins with Tops>20 in at least one of the baits. Norm Correlation was used as a distance metric with 2 filter functions: Neighborhood lens 1 and Neighborhood lens2. Resolution 30 and gain 3 were used here using the Ayasdi 3.0 Cure 3.0 software ^24^. Proteins are colored based on the rows per node. Color bar: red: high values, blue: low values. Node size is proportional with the number of proteins in the node. (B) A reduced network was created using information from the CRApome database, and the network was generated using Cytoscape platform ^23^. Proteins associated with CBX1, CBX3, and CBX5 are expanded and highlighted in the inset. (C) Hierarchical biclustering using PermutMatrix ^40^ of protein with extreme positive TopS values. The 28 proteins with the highest TopS values across the dataset are shown. Rows and columns were clustered using Pearson as the distance and Ward linkage as the method. Yellow corresponds to high TopS values and grey shows negative TopS values. Proteins that are not present in the purifications are in black.

There are many additional approaches to use to further interpret the dataset. For example, we introduced another filter to the TopS associations by adding information from the CrapOme, which is a database of proteins known to be present in the 411 negative controls ^27^. Here, we selected proteins with high TopS values in our dataset that are present in a maximum of 10/411 controls with a maximum spectral count of 15. We created a reduced network and inspected these specific interactions (Fig. 3B). For simplicity, we focused on the interactions of the CBX proteins where we observed a high connection between a set of zinc finger proteins with KRAB domain and the CBX proteins (Fig. 3B). Note that most of these zinc finger proteins are not identified in any of the 411 negative controls, and are likely specific to the current dataset. KRAB zinc finger proteins play an important role in the evolution of gene regulatory networks ^28^, and one KRAB protein named TRIM28/KAP1 has been shown to directly interact with HP1 proteins (CBX1, 3, and 5) ^29, 30^. The CBX1, 3, and 5 subnetwork presented in Figure 3B suggests a large number of additional KRAB Zinc finger proteins associate with CBX proteins in unexplored processes.

The known direct interactions of HP1 proteins (CBX1, 3, and 5) with TRIM28 ^29, 30^ suggested we look deeper into TopS values in our dataset. We sorted the TopS values for all the prey proteins for each bait protein (Supplemental Table 2) from highest to lowest to observe the extreme negative and positive values in the dataset. The 28 prey proteins with the highest TopS values across the 17 human DNA repair bait purifications are shown in Figure 3C. TRIM28 is the highest scoring protein with all three HP1 proteins (CBX1, 3, and 5). In every other bait, except RPA3, TRIM28 is amongst the ten most negative TopS values and TRIM28 is the most negative scoring protein in the CBX8, MSH6, and WDR76 baits (Supplemental Table 2). This pattern of very high TopS values in certain baits being extreme negative values elsewhere in the dataset occurs for several additional bait:prey interactions. This includes XRCC5 and XRCC6, which also directly interact to form a heterodimer ^13^. In the XRCC5 affinity purification, XRCC6 is the highest scoring prey protein (Supplementary Fig. 2), which is the highest TopS value in the entire dataset, and in the XRCC6 affinity purification XRCC5 is the highest scoring prey protein (Fig 3C and Supplemental Table 2). XRCC5 and XRCC6 are amongst the most negative TopS values in several other bait purifications including CBX5, CBX8, MSH2, MSH3, MSH6, RPA2, RPA3, SPIN1, and WDR76 (Fig 3C and Supplemental Table 2). These results suggest that extremely positive TopS values reflect direct protein:protein interactions.

As an example of how to further utilize prey proteins with high TopS values, we next sought to investigate the therapeutic aspect of these interactions since proteins involved in DNA repair pathways are known for their role in human diseases like cancer. We used WebGestalt ^31^ to perform drug-gene association enrichment for the proteins with high TopS scores in our dataset, and the enrichment analysis resulted in 9 identified drug classes (Fig. 4A).

**Figure 4.**
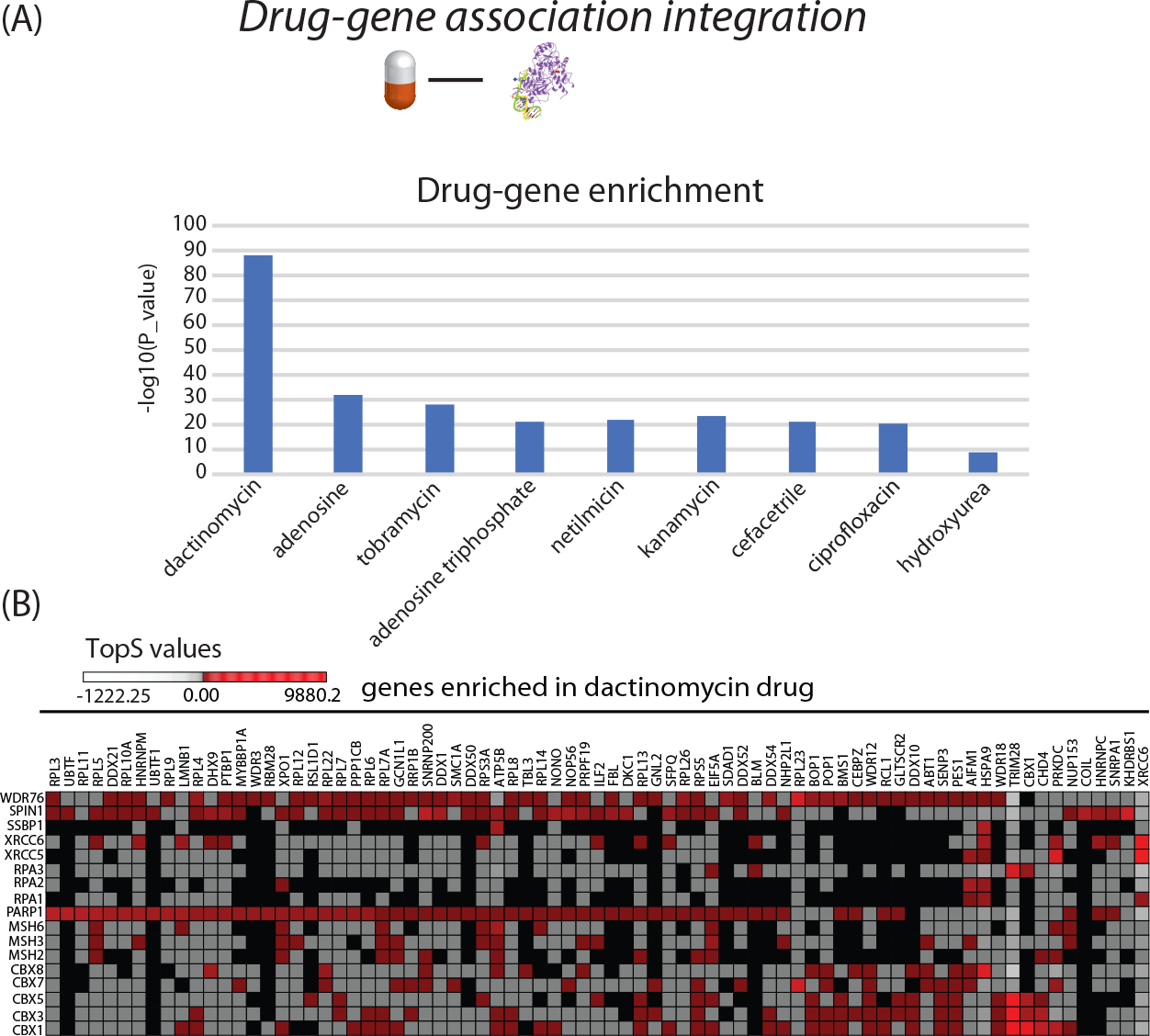
Enrichment of Drug Targets in the Human DNA Repair Network. (A) The proteins detected and scored in our human DNA repair dataset were enriched in 9 gene-drugs classes. The enriched classes were obtained using WebGestalt database ^31^. (B) Proteins/Genes enriched in the dactinomycin set are represented. The red color represent high TopS values.

Unexpectedly, we observed that the enriched classes consist of proteins with high topological scores in the same bait (Supplementary Table 5). For example, the association with dactinomycin is enriched with most proteins with high positive TopS scores in the PARP1 and WDR76 baits (Fig. 4B and Supplementary Table 5), whereas tobramycin for instance is enriched with proteins with high scores in SPIN1 (Supplementary Table 5). Dactinomycin is approved for use in treatment of several cancers, like Wilms’s tumor ^32^. These enrichment results indicate that TopS can highlight interactions that are targeted by drugs in a dataset.

### TopS Analysis of Yeast Chromatin Remodeling Complexes

The fact that the highest TopS value in the human dataset was XRCC6 in the XRCC5 affinity purification, which is a known heterodimer ^13^, led us to investigate the TopS values in a well-characterized yeast chromatin remodeling system. We have previously utilized deletion network analyses to determine the modularity of the INO80 ^14^ and SWI/SNF ^12^ chromatin remodeling complexes. Here, we reanalyzed two published yeast INO80 ^14^ and SWI/SNF ^12^ protein complexes datasets for which cross linking data also exists ^33, 34^. However, we only considered the wild type affinity purifications and did not consider the affinity purifications in genetic deletion backgrounds. We sought to determine if TopS analysis applied to the wild type INO80 and SWI/SNF data could capture modules and potential direct interactions as determined from cross linking data.

Identifying direct interactions from quantitative wild-types affinity purification datasets has been a long-standing problem. Examining the proteins abundances in separate samples, one cannot identify the direct interactions using standard approaches. This challenge is illustrated in Figure 5 where the protein abundances, as estimated by dNSAF, of the INO80 complex in ARP8 bait (Fig. 5A) and the protein abundance of the SWI/SNF complex in the SWP82 bait (Fig. 5B) are shown. For example, ACT1, ARP4, ARP8, IES4, and TAF14 are known to be part of a module based on deletion network analysis ^14^ and cross linking data ^34^, yet their dNSAF abundances have no particular pattern.

**Figure 5.**
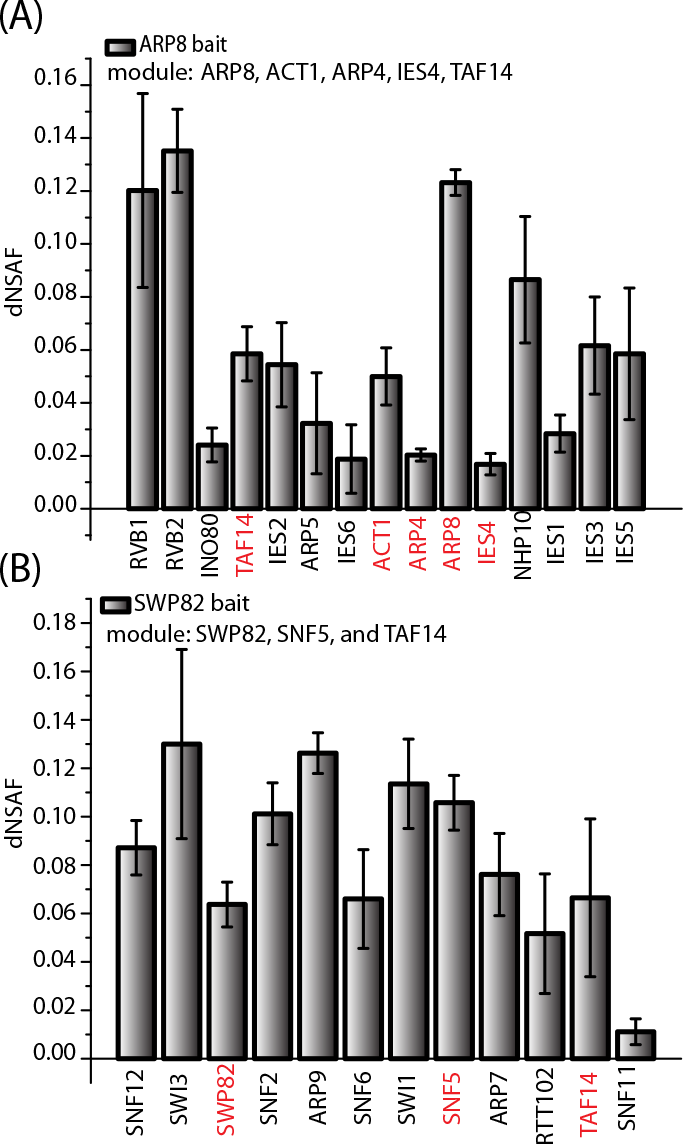
Failure to Capture Modularity in Wild Type Yeast Chromatin Remodeling Complexes with a Standard Approach. Distributed normalized spectral abundance factors for the subunits of the INO80 and SWI/SNF complexes in APMS experiments using ARP8 and SWP82 as baits. In red are colored proteins included in (A) the ARP8, ACT1, ARP4, IEAS4, and TAF14 module, and (B) the SWP82, SNF5, and TAF14 module.

To determine whether TopS values can capture the modularity and direct protein interaction in these complexes, we merged a total of 24 wild type affinity purifications from INO80 and SWI/SNF (Supplementary Tables 6 and 7). First, the INO80 data ^14^ was preprocessed by extracting the non-specific proteins, and proteins that passed the contaminant extractions in INO80 purifications (Supplementary Table 6) were also searched in the SWI/SNF dataset (Supplementary Table 7). A total of 237 proteins were shared between the two complexes and included in a merged dataset. TopS analysis was applied to both complexes and the topological scores are reported in the Supplementary Tables 6 and 7. As with the human DNA repair network, here we directly imported TopS values into a TDA based network analysis (Figure 6).

**Figure 6.**
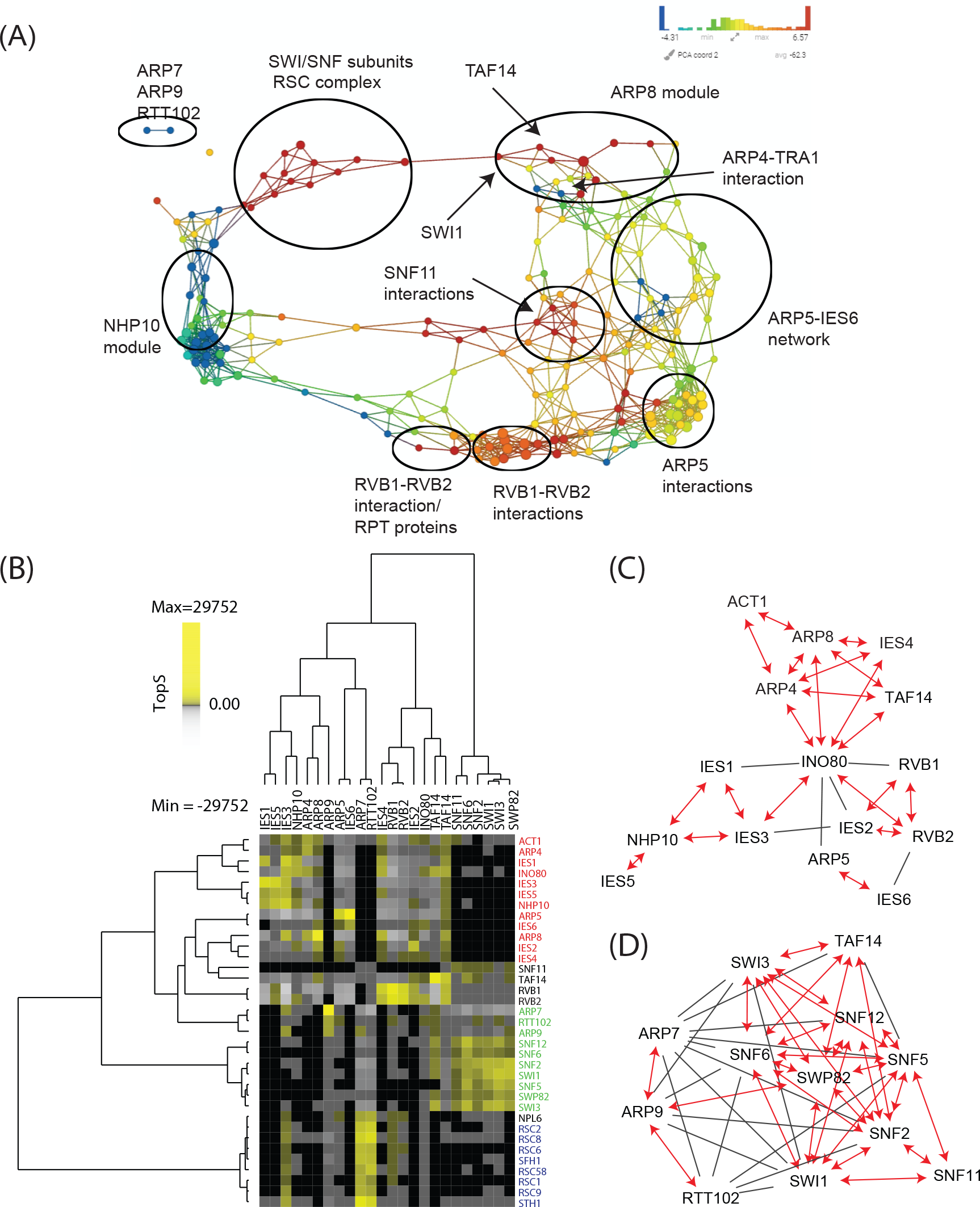
Topological Network and Comparative Linkage Analysis of Yeast Chromatin Remodeling Network. (A) A TDA network was constructed for the yeast data inputting TopS scores to the Ayasdi 3.0 Cure 3.0 software ^24^. Distance Correlation was used as a metric with 2 filter functions: Neighborhood lens 1 and Neighborhood lens2. Resolution 50 and gain 6x eq. were used. Proteins are colored based on the rows per node. Color bar: red: high values, blue: low values. Node size is proportional with the number of proteins in the node. The location of protein complexes subunits within the topological network are drawn on top of the TDA output. (B) Hierarchical biclustering using TopS values. Components of the INO80, SWI/SNF, and RSC complexes were clustered using PermutMatrix ^40^. Rows and columns were clustered using Pearson as the distance and Ward linkage as the method. Yellow corresponds to high TopS values and grey shows negative TopS values. Proteins that are not present in the purifications are in black. (C)-(D) Comparison of TopS values with crosslinking data. Each plot shows the direct interactions between subunits of (C) INO80 and (D) SWI/SNF. A bidirectional edge and red color represents the reported crosslinking interactions with a high TopS value. In black, we represent the reported crosslinking interactions with lower TopS values.

TopS values of wild type affinity purifications were able to identify the modules of the INO80 and SWI/SNF complexes (Fig 6A) comparable to the results from deletion network analyses ^12, 14^. For example, in the INO80 complex, the ARP8 module is separated from the NHP10 module and from ARP5-ARP6 module (Fig. 6A, Supplementary Fig. 3). Interestingly, the ARP8 module is connected to a module of the SWI/SNF complex since both modules share the TAF14 protein (Fig. 6A). In addition, the ARP9, ARP7 and RTT102 module in SWI/SNF was also clearly separated from the other modules (Fig. 6A). Lastly, the RSC complex is present in the dataset, especially in the ARP7 and RTT102 affinity purifications, and ARP7, ARP9, and RTT102 are part of both the SWI/SNF ^12^ and RSC ^35^ complexes. Additional components of RSC are present in a SWI/SNF subunit/RSC complex portion of the network (Fig. 6A).

Next, similar to the human DNA repair network, we sorted the TopS values for all the prey proteins for each bait protein (Supplemental Tables 6 and 7) from highest to lowest to observe the extreme negative and positive values in the dataset. The 35 prey proteins with the highest TopS values across the 24 yeast chromatin remodeling wild type bait purifications are shown in Figure 6B. The highest TopS value in the entire dataset is ARP5 in the IE6 affinity purification. Furthermore, ARP5 and IES6 are the only two proteins in the IES6 affinity purification from the INO80 complex with positive TopS values, all the other proteins in the INO80 complex have negative values in the IES6 affinity purification (Fig. 6B and Supplemental Tables 6 and 7). ARP5 and IES6 are well-characterized direct interactors in a sub complex within the INO80 complex ^14, 36^. A similar result occurred in the ARP9 affinity purification where ARP7, RTT102, and ARP9 had the three highest TopS values, and all the other components of the SWI/SNF and RSC complexes had negative values in the ARP9 affinity purification (Fig. 6B and Supplemental Tables 6 and 7). Again, ARP7, RTT102, and ARP9 are a known module since they are shared by both the SWI/SNF ^12^ and RSC ^35^ complexes.

Lastly, we next evaluated the overlap of our high TopS values with reported crosslinking interactions from INO80 ^34^ and SWI/SNF ^33^ (Fig. 6C-D and Supplementary Tables 6 and 7). Our results showed a high overlap between crosslink interactions and proteins-baits pairs with high TopS values. For the INO80 complex, we observed 77% overlap, while for the SWI/SNF complex 63% overlap was observed. The direct interactions are mostly between subunits located within the same module and TopS identified the majority of these interactions (Fig. 6 C-D). The major exceptions are for proteins that are shared between different complexes like RVB1, RVB2, ARP7, ARP9, and RTT102 (Supplementary tables 6 and 7). In these cases, TopS give the highest scores to other complexes which share these proteins. For example, the module ARP7, ARP9, and RTT102 is shared with the RSC complex and we can see from Fig. 6B, that members of the RSC complex are highly enriched in the ARP7 purifications. Overall, we found that TopS values from an analysis of wild type INO80 and SWI/SNF APMS captured known modularity and direct interactions from these protein complexes.

## Discussion

To predict protein interactions from affinity purifications using quantitative proteomics, we have devised a new topological scoring approach for evaluating the preference for each protein in a bait relative to other baits. We have built on the concepts of topological data analysis ^24^ as applied to protein interaction network analysis ^14, 25^ to devise the TopS approach. Here, we combine information from row, column, and total distributed spectral counts into this new score to differentiate the preference of interaction with baits. This is specifically important for cases where proteins are detected in many runs. We illustrate the methodology and its advantages through the analysis of a human DNA repair protein interaction network generated using transient transfections and a yeast chromatin remodeling protein interaction network dataset. TopS values were directly incorporated into clustering and network assembly approaches and provided unique insights into both protein interaction networks.

TopS values cover a broad and meaningful negative to positive range. In the human DNA repair protein interaction network dataset, the highest TopS value was XRCC6 in the XRCC5 affinity purification and these two proteins are known to interact and form a heterodimer ^13^. Furthermore, XRCC5 and XRCC6 are amongst the most negative TopS values in several other bait purifications in the human DNA repair network dataset. The highest TopS value in the entire dataset is ARP5 in the IE6 affinity purification, and again these two proteins are a well-characterized, directly interacting, submodule of the INO80 complex ^14, 36^. Again, ARP5 and IES6 had negative values in other INO80 bait purifications. Extreme positive TopS values in specific baits are typically reflected as extreme negative values in other bait purifications, even if the bait protein is part of the same complex. This distinguishing feature of TopS provides unique insights into proteins within a complex and suggests the presence of direct interactions. Furthermore, in the INO80 and SWI/SNF analysis, we were able to capture modularity from wild type affinity purifications that we previously could only capture with the analysis of affinity purifications in deletion mutant backgrounds ^12, 14^ and our results correlated strongly with crosslinking data ^33, 34^. A second important feature of TopS is that it is hence capable of capturing modules within wild type protein complex datasets.

The TopS platform has several important features. It is easy to implement. There are no parameters or assumptions of a probability distribution in our algorithm. The number of replicates can vary for the baits, where the replicates could be averaged out by the user, without severely affecting the results. TopS values can serve as a starting point or as a scaffold for other computational methods, visualization tools, and can be directly used in clustering and network analysis approaches. TopS can be extended to other quantitative values, such as peptides, ion intensity, or percentage post translational modification, and can be applied to data from any species. Lastly, multiple datasets can be integrated by merging their TopS scores.

For straightforward usage by the community, TopS is implemented as a SHINY application (https://shiny.rstudio.com/) and available as supplemental material. TopS is complementary to many computational pipelines where quantitative proteomic analysis of affinity purifications is used. Approaches like COMPASS ^5^, QSPEC ^6^, SAINT ^7^, and SFINX ^8^ largely yield statistical values, like a *p* value, to provide a confidence that two proteins are associating or are part of a protein complex. The TopS platform can be easily implemented in addition to these approaches to further analyze protein interaction network datasets. If quantitative values are provided for all the prey proteins in a given bait, TopS can be used to reanalyze data, like the recent description of the human polycomb complexome ^4^, for potential direct interactions and/or modules within protein complexes.

## Methods

### Materials

Magne™ HaloTag^®^ magnetic affinity beads were purchased from Promega (Madison, WI). The following clones from the Kazusa DNA Research Institute (Kisarazu, Chiba, Japan) were used: Halo-WDR76 (FHC25370), Halo-XRCC5 (FHC07775), Halo-XRCC6 (FHC01518), Halo-RPA1 (FHC01462), Halo-RPA2 (FHC11655), Halo-RPA3 (FHC06678), Halo-MSH2 (FHC07773), Halo-MSH3 (FHC12698), Halo-MSH6 (FHC08173), Halo-CBX1 (FHC07438), Halo-CBX3 (FHC02188), Halo-CBX5 (FHC10519), Halo-CBX7 (FHC10535), Halo-CBX8 (FHC01705), Halo-PARP1 (FHC01012), Halo-SPIN1 (FHC10419), and Halo-SSBP1 (FHC07926).

### Affinity Purification and Quantitative Proteomic Analysis

Human proteins in the pFN21A plasmid with an N-terminal HaloTag™ were transiently transfected into HEK293T cells, whole cell extracts prepared and Halo™ affinity chromatography was performed on each independent whole cell lysate as described previously^37^. Samples were processed and analyzed using label free quantitative proteomic analyses as described previously ^19^. Briefly, TCA-precipitation, LysC/Trypsin digestion, and multidimensional protein identification technology (MudPIT) on linear ion trap mass spectrometers (LTQ, Thermo Scientific) were performed as previously described ^19^. RAW files were converted to the ms2 format using RAWDistiller v. 1.0, an in-house developed software. The ms2 files were subjected to database searching using SEQUEST (version 27 (rev.9)) ^19^. Tandem mass spectra of proteins purified were compared against 29,375 non-redundant human protein sequences obtained from the National Center for Biotechnology (2012-08-27 release). Randomized versions of each non-redundant protein entry were included in the databases to estimate the false discovery rates (FDR) ^19^. All SEQUEST searches were performed without peptide end requirements and with a static modification of +57 Daltons added to cysteine residues to account for carboxamidomethylation, and dynamic searches of +16 Daltons for oxidized methionine. Spectra/peptide matches were filtered using DTASelect/CONTRAST ^38^. In this dataset, spectrum/peptide matches only passed filtering if they were at least 7 amino acids in length and fully tryptic. The DeltCn was required to be at least 0.08, with minimum XCorr value of 1.8 for singly-, 2.0 for doubly-, and 3.0 for triply-charged spectra, and a maximum Sp rank of 10. Proteins that were subsets of others were removed using the parsimony option in DTASelect on the proteins detected after merging all runs. Proteins that were identified by the same set of peptides (including at least one peptide unique to such protein group to distinguish between isoforms) were grouped together, and one accession number was arbitrarily considered as representative of each protein group. Quantitation was performed using label-free spectral counting. The number of spectra identified for each protein was used for calculating the distributed normalized spectral abundance factors (dNSAF) ^21^. NSAF v7 (an in-house developed software) was used to create the final report on all non-redundant proteins detected across the different runs, estimate false discovery rates (FDR), and calculate their respective distributed Normalized Spectral Abundance Factor (dNSAF) values. All the DNA repair data used in here are deposited at the https://massive.ucsd.edu/ with MassIVE ID # MSV000081377. The data used for the analysis of the S. cerevisiae INO80 and SWI/SNF chromatin remodeling complexes are from previous studies on INO80 ^14^ and SWI/SNF ^12^

### Model for predicting interaction preferences

#### Normalization method for overexpressed baits

Since our approach aims to identify the enrichment of each protein in each bait relative to a collection of baits, overexpression of affinity-tagged bait proteins can diminish the interaction score. With this knowledge, we therefore selected to use a normalization method where the baits are estimated directly from the dataset.

To adjust for baits enrichment, we used this approach

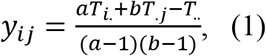

where a is the number of columns, b is number of rows, T_i_ represents the bait total spectral counts, T_j_ is the row total spectral counts, and T. is the total spectral counts in the matrix. Using this approach, the estimated values of the bait proteins are now close to their average spectral counts in all the APMS runs of the dataset.

#### Calculation of topological scores

All the data that passed criteria from the QSPEC analysis was used as an input to the TopS determinations.

We used a simple model to calculate a score for each prey bait interaction as follows:

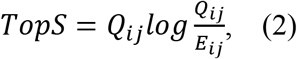

where Qij is the observed spectral count in row i and column j; and

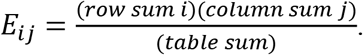

The data is treated in the following manner: 1) For each column and row, the sum of spectral counts is calculated; 2) the total number of spectral counts in the dataset is determined; 3) a TopS value for each protein in a bait APMS run is determined as described in the equation (2). If the actual number of spectra of a prey protein in a bait APMS run exceeds that in all APMS results being analyzed, the presumption is that we have a positive interaction preference. Likewise, fewer spectra than in the APMS runs indicate a negative interaction preference. Proteins that were not detected in a particular run were assigned the value 0. A high positive score indicates a direct interaction between bait and prey. A negative score suggests that prey protein and bait interact to form a complex elsewhere in the dataset. We applied this framework to construct signed interaction networks derived from two independent APMS datasets obtained for yeast chromatin remodelers and human DNA repair proteins.

#### Implementation

TopS is written using Shiny application (R package version 3.4.2) for R statistics software. TopS uses several packages, including gplots, devtools and gridExtra. TopS is freely available and packaged as a compressed archive, in a *.zip format, and users can download the compressed file to their machine and decompress the file.

#### Data input

TopS take a numeric data matrix as input where multiple dimensions (e.g. proteins) are measured in multiple observations (e.g. baits/samples). In our case, the numerical values are distributed spectral counts. To make data input easier for the end user, we have defined the input file formats that include rows and columns annotation and numeric data. Any quantitative value, not only spectral counts, can therefore be utilized. TopS include an example dataset for testing purposes: DNA repair data set.

#### Data output

TopS generates an automatic output to make this type of analysis easier. *MS Excel* can be used to visualize the output and identify for example differences between samples or interactions between proteins and baits. Pearson correlation map, clustering analyses on initial numeric values (in our case distributed spectral counts) and TopS values are provided to the user as a pdf format output.

### Topological data analysis (TDA)

The input data for TDA are represented in a bait–prey matrix, with each column corresponding to purification of a bait protein and each row corresponding to a prey protein: values are TopS values for each protein. A network of nodes with edges between them is then created using the TDA approach based on Ayasdi 3.0 Cure 3.0 software (AYASDI Inc., Menlo Park CA) ^24^ as described previously ^14^. Here, Norm Correlation was used as a distance metric with 2 filter functions: Neighborhood lens 1 and Neighborhood lens2. Resolution 30 and gain 3 were used to generate Fig. 3A. and Resolution 50 and gain 6x eq. were used to generate Fig. 6A.

## Acknowledgements

Research reported in this publication was supported by the Stowers Institute for Medical Research and the National Institute of General Medical Sciences of the National Institutes of Health under Award Number RO1GM112639 to MPW. The content is solely the responsibility of the authors and does not necessarily represent the official views of the National Institutes of Health.

## Author Contributions

M.E.S. and M.P.W. designed the experiments. J.M.G., B.D.G., and A.D. performed experiments. M.E.S. performed computational analyses of data. M.E.S, L.F., and M.P.W. wrote the manuscript.

## Competing Interests

The authors declare no competing financial interests.

